# Gene Expression and Physiological traits in Mice

**DOI:** 10.1101/2023.05.30.542939

**Authors:** Rafael Ramos, Igor Nelson

**Affiliations:** University of Coimbra, 2022129995; University of Coimbra, 200911124

**Keywords:** Big Data Analysis, Statistical Tests, Health Indicators

## Abstract

**Background:** Gene expression regulates several complex traits observed. In this study, datasets comprising of transcriptome information and clinical traits regarding fat composition and vitals were analyzed via several statistical methods in order to find relations between genes and clinical outcomes.

**Results:** Biological big data is diverse and numerous, which makes for a complex case study and difficulties to stablish a metric. Histological data with semi-quantitative scores proved unreliable to correlate with other vitals, such as cholesterol composition, which complicates prediction of clinical outcomes. A composition of vitals, turned out to be a better variable for regression and factors for gene analysis. Several genes were found to be statistically significant after statistical analysis by ANOVA regarding the progressive categories of the preferred clinical variable.

**Conclusions:** ANOVA is proposed as a method for genetic information retrieval in order to extract biological meaning from RNA seq or microarray data, accounting for multiple classes of target variables. It Provides a reliable statistical method to associate genes or clusters of genes with particular traits.

**Supplementary information:** Supplementary data are available in annexes.

## 1 Introduction

The relevance of statistics in biology is a longstanding one, with great statisticians being also great biologists[1]. Nowadays it is arguably even more so, with the increasingly accessible computation power and highly parallelized assays that revolutionized biology, leading to a paradigm shift from hypothesis-driven research to data driven research[6]. A more in-depth discussion on the theme can be found on the Special Issue of Studies in History and Philosophy of Science Part C: Studies in History and Philosophy of Biological and Biomedical Sciences entitled Data-Driven Research in the Biological and Biomedical Sciences. An accurate understanding of statistical tools is central to a proper research and it’s misuse maybe central in the reproducibility crisis facing biomedical research[7,8]. The broad objective of this article is first and foremost the use of statistical tools to analyse a biological dataset, regarding in depth the assumptions required for every step.

The advances in biology the made possible high throughput screening and data collection of various biological molecules gave rise to the “omics” science, i.e. the large scale study of most if not all biomolecules in a sample. It is therefore important to emphasize the central role that statistics play in all steps of these studies as it is through statistical methods that one can derive conclusions from the data. A deeper understanding of how genomic expression can affect certain phenotype or lead to diseases or worse prognostics, made possible through statistics, is a major goal not only for research, but as guidelines for decision makers and public policies.

One such public health concern is obesity. In that sense, many studies are performed to find underlying causes or risk factors for development and prevalence of obesity [9]. In 2006, Ghazalpour et al. [2] studied the relation between genes and weight on mouses. In their work, a crossbreed, named BhX, was subject to a Western diet and had their clinical traits evaluated, such as body composition and biomarkers, and had their livers subjected to microarray analysis, generating a rich dataset.

Their approach to this analysis was the Weighted Gene Correlation Network Analysis (WGCNA)[5] in order to find clusters of coexpressed genes and the relation of clusters of these genes to clinical outcomes.

While Pearson’s correlation coefficient is intimately related to statistical properties and regression analysis, it does very little for statistical hypothesis testing. Since this article’s aim is to use an array of statistical methods for data exploration and hypothesis testing and it cannot therefore follow the described protocol. However, the dataset generated is still rich with various types of variables, categorical and continuous, and provide good material for statistical analysis with significant biological conclusions.

## 2 Material and Methods

The datasets used for this article were retrieved from WGCNA’s tutorial hosted at https://horvath.genet-ics.ucla.edu/html/CoexpressionNetwork/Rpack-ages/WGCNA/Tutorials/. The description on how the dataset was generated is better found on Ghazalpour, et al. 2006[2]. The exploration and statistical analysis of the dataset was performed in R, as will be explained in depth further. The package caret[4] was used to clean and process data.

### 2.1 Data Exploration

The datasets available for analysis are separated into four files, two are gene expression datasets for male and female mice, one has clinical information regarding each mouse and the last file has gene annotations. Both genomic datasets had only 3600 of the most variant probes, all standardized and with few missing values, so little had to be done regarding cleaning these datasets genewise. The samples on the other hand were grouped via hierarchical clustering with an Unweighted Pair Group Method with Arithmetic mean (UPGMA) and extreme outliers (> *Q*_3_ + 3 × *IQR*) were removed.

The clinical dataset had greater diversity of data types. Qualitative variables, such as *sex*, were processed into suitable factors. Continuous variables had their distribution analysed to check for normality and extreme outliers. Logarithmic and square root transformations were applied to some of those variables in order to normalize the distribution. The remaining variables were normalized via Box Cox transform. Extreme outliers were removed by the same method described before. Date variables were combined as start and endpoint, turning into counter of days. The normalized continuous variables were then centered and scaled. Variance and correlation were assessed, variables with near zero variance and correlation above 0.9 were removed. Also, a new variable on metabolic syndrome was creating compressing information on abdominal fat, triglycerides, glucose, insulin and HDL cholesterol was created.

To have a general idea of a dataset with that many variables it is important to find ways which the information can be preserved and represented. In that sense techniques of dimensionality reduction come in handy. In this case, PCA was done using caret.

### 2.2 Statistical Analysis

The clinical dataset was mainly analyzed regarding the relation between variables mainly using being scatterplots and linear regression models. A few select variables were subject to stepwise regression in order to check their fit for interesting variable for disease prediction, such as aortic lesions. The assumptions and quality checks of those regressions were individually assayed, i.e., the predicted values have a linear relationship with real values. the distribution of residues being normal, homoscedastic, with no outliers, and no leverage values, which can be indirectly inferred with Cook’s distance.

Two random variables are independent if the realization of one does not affect the probability distribution of the other. Since the Gender factor(given by *sex* variable in the dataset) has only male and female levels, accounting for differences in a measurement can be seen as a statistical test to check for difference between two populations, regarded the proper test assumptions.

Considering the aim of this study is to check for differences in mouse weight, it follows that mice should be separated into weight categories. To that end, they were separated in a factor with four levels, for lean, normal, obese, and very obese. In that sense, MANOVAs and ANOVAs provide a powerful tool for assaying the relevance of weight in the outcome of the different vitals, provided, once more, the assumptions are met.

On account of the gene expression being already separated in two tables, only the female expression was analyzed. Due to the continuous and normally distributed nature of the interesting clinical variable, defined by the regression metrics and characteristics, it was split into quantiles of increasing intensity, from which a factor of four levels was derived. This factor was used as independent variable for an ANOVA, to test is a gene differs from light to intense cases of metabolic syndrome. This was done for all genes, and the assumptions of normality and homoscedasticity were checked via Shapiro and Levene’s test. The genes that passed the assumptions (regardless of the p-value) were subjected to a *post hoc* Tukey HSD (Honest Significant Difference) test, with Bonferroni’s correction of p-value, to find the pairwise differential expression between different levels. The Lowest vs Highest comparison was then subject to Benjamini-Hochberg[10] adjustment of p-value and the differentially expressed genes were shown via volcano plot.

Identifying the functions of genes was beyond the scope of this article, but they were able to be identified by the gene annotation table of the dataset.

## 3 Results

Data cleanup is an essential part of the process, and has to be regarded first, being it for the understanding of what are the relevant features to be analyzed, or to ensure that they are well behaving regarding the statistics.

Once they are properly treated, statistical inquiries may begin. It is important to note, however that even in normalized distributions, some assumptions may or may not hold, as we will see further on that, even a simple comparison of two normalized population distributions of cholesterol cannot be done by a Pooled t-test and must be done as a Welch’s t-test.

The statistical analysis on the clinical dataset is central to a better understanding of biological phenomena and has an easier biological interpretations than expression data. Therefore, a good analysis on the composition and relation of clinical attributes of samples is relevant not only to a better understanding of the biology as to indication on how this information can be extrapolated to analyze gene expression and molecular function.

So, in this sense, biological and applied biostatistical research is three-fold: first, the understanding of the dataset leads to the perception of possible analytical paths; the understanding of the experimental or clinical factors, subject to some analytical path, which leads to the perception of the biological phenomena; and the analysis of the composition of samples, which leads to the understanding of the molecular mechanisms relevant to the biological outcome. In this spirit, the results will be presented following the same order.

### 2.1 Dataset exploration

The clinical dataset consists of 361 observations of 37 variables. Of those, the categorical variables *Strain* and *Sex* were selected. Numerical variables regarding weight, length, abdominal fat, other fats, total fat, triglycerides, total cholesterol, HDL cholesterol, Free fatty acids, LDL and VLDL, MCP-1, Glucose, Insulin, Glucose x Insulin ratio, Leptin, Adiponectin and Aortic Lesions were kept for the analysis. All these variables have objective composition units such as μg/mL, except Aortic Lesions, which is scored semi quantitatively using histological information.

The dates of sacrifice and beginning of diet were used to calculate the amount of time spent in the treatment. Save for few outliers (<50 and >140) the mice spent between 100 and 125 days in a Western diet. However, after some exploration it seems this amount of time is not relevant for clinical outcomes.

Another such variable that was observed, yet not included in analysis due to sample size and lack of genetic information was Strain. Only the BhX crossbreed, also regarded as F2, was subject to microarray analysis. However, there is still clinical data regarding the parents (F1) and the original strains, C57BL/6 (B6), a susceptible strain to obesity, and C3H a resistant one. One can see the effect of this weight distribution via the violin plot, seen in Figure 1, which is an interesting result.

**Figure 1:**
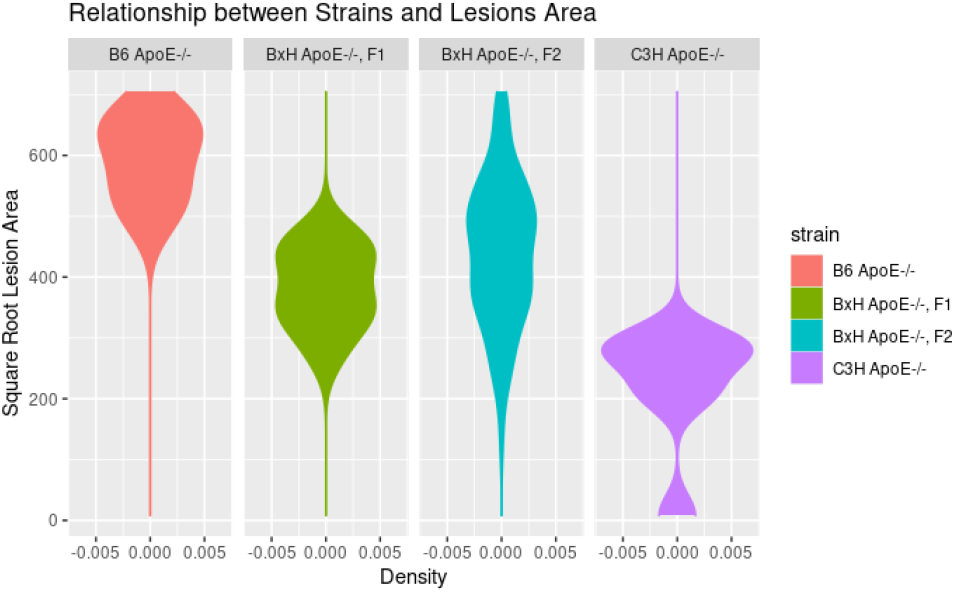
Violin plots of aortic lesion area regarding strain. The susceptible b6 strain has the highest values, while the resistant C3H the lowest. F2 crossbreed has a much more variate phenotype.

From the clinical dataset, Triglycerides and MCP-1 were normalized after a log transform and Leptin and Adiponectin were normalized by square root. Insulin and Glucose/Insulin were subject to BoxCox. While HDL was mostly normal, there was a very extreme outlier that interfered with each and every measurement regarding this variable and was subsequently removed. After checking complete cases 279 samples remained.

There were no variables with near zero variance, but Total cholesterol and Insulin were highly correlated with LDL+VLDL and glucose/insulin, respectively.

The expression datasets had each 3600 genes, with 132 male samples and 143 female samples. After cleaning the dataset some samples were dropped, resulting in the loss of clinical information for that sample and subsequent removal of the expression dataset. The working tables had 3600 for 100 male samples and 110 female samples.

Lastly, a new variable called metabolic syndrome was created by taking the mean of abdominal fat and triglycerides, discounted the influence of HDL cholesterol and glucose/insulin.

Considering the amount variables, a simple plot cannot convey all the information available in the dataset. To circumvent this, one can use techniques for dimensionality reduction. In this dataset, once normalized so no variable overpowers others, a PCA was executed represented in Figure 2.

**Figure 2:**
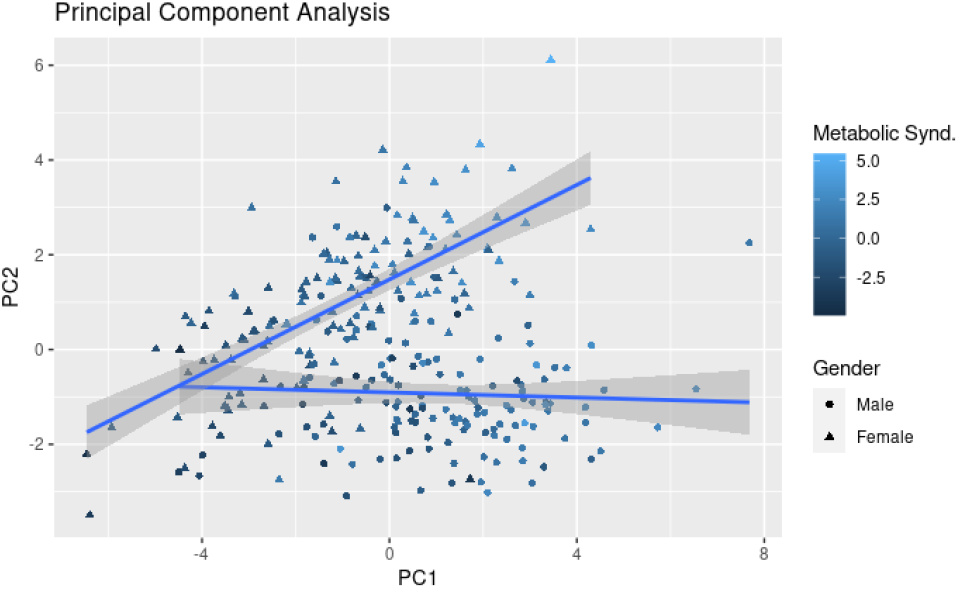
PCA analysis of clinical information dataset. Color scheme from dark blue to light blue varies according to the metabolic syndrome variable, while shapes relates to sex, dots for male, triangles for female. Regression shows different gender behavior.

### 2.2 Clinical traits

Clinical traits were evaluated firstly as scatterplots between highly correlated variables, such as sugars and triglycerides as can be seen on Figure 3, or weight and fat. While the relationship is mostly linear, when accounting the abdominal fat against weight there are two groups, as seen in Figure 3. While the relationship wasn’t immediately obvious at first glance, when accounting for the difference in sex, the difference in metabolic processes and responses to fat accumulation becomes clear. Female mice have a very linear response to weight gain as abdominal fat, meanwhile male mice don’t have that clear a relationship, operating at around a mean value of abdominal fat, mostly regardless of weight distribution.

**Figure 3:**
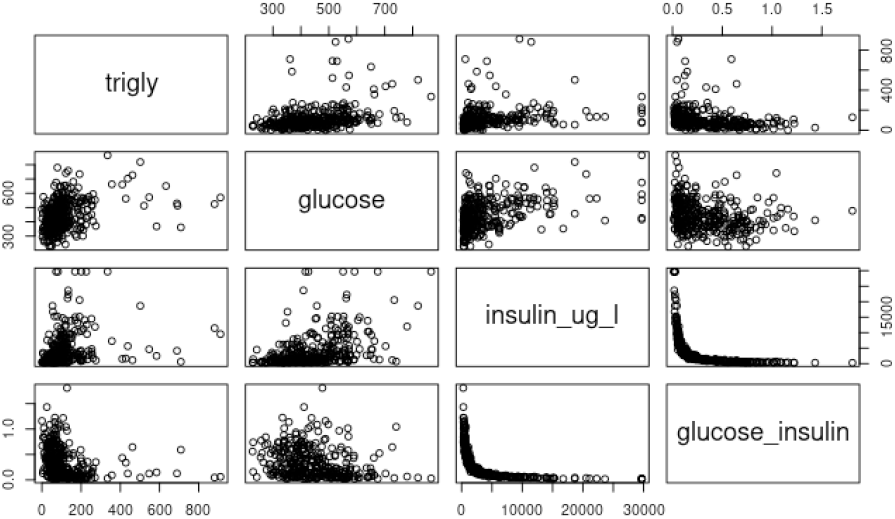
Scatterplots between different variables. Correlation and linear dependency can be inferred and observed. It is important to note that insulin and glucose by insulin have a nonlinear relationship.

Next the linear relationship between variables relating to the outcome of the condition, i.e. aortic lesions and metabolic syndrome. To that end, two multilinear models for prediction of were made, then improved by stepwise regression, one for each dependent variable.

Multiple linear regression of aortic lesions showed p-values for most variables being not significant, and an overall R^2^ of 0.225. The most important variables for the multilinear model were the strain and triglycerides. The relationship between lesions and strain has already been graphically seen on Figure 1.

The Akaike information criterion (AIC) is a mathematical method for evaluating how well a model fits the data it was generated from. Since so many variables have a small influence on the model, the overall best model concerning AIC had strain, triglycerides, adiponectin, insulin and glucose /insulin. It, however still performed poorly, with an R^2^ of 0.233.

When performing a linear model, there are many metrics beside R^2^ that one must be wary, mainly concerning the residuals. The residual distribution should be normal, homoscedastic and with no outliers, the predicted and observed values should follow a linear relationship and there should be little to no leverage observations. Cooks distance, which is a metric describing the impact the removal of an observation has on the overall model, should be lower than one for all observations. Plots regarding this quality measures for the stepwise regression of aortic lesions can be found on supplementary information.

As observed in Figure 4, male and female mice have different behaviors. To assess the difference in a more objective way, a statistical test was performed regarding the abdominal fat. As mentioned before, the statistical assumptions of tests should be met to extract meaningful conclusions. From Figure 5 is possible to see that the distributions seem normal. While there are outliers, none are extreme. The independence of observations is derived from the experiment conditions. Lastly, however is the homoscedasticity condition, which one can infer from Figure 5, is not met. As such, one must use the Welsh test due to different variances. The results of the test (p-value < 2.2e-16) indicate it is safe to reject the null hypothesis and say there is significant difference on the abdominal fat content between male and female mice.

**Figure 4:**
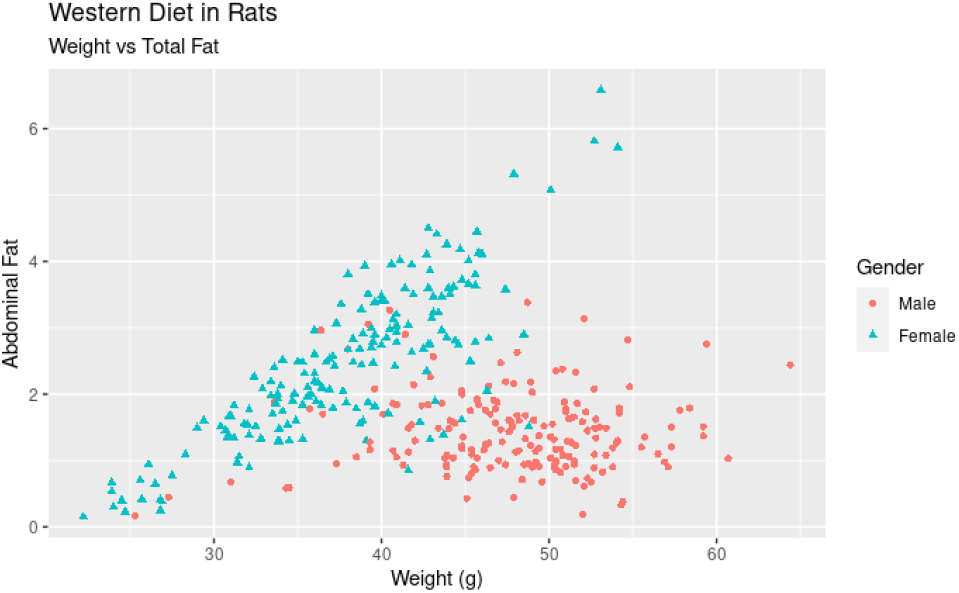
Scatterplot of Abdominal Fat vs Weight. The two different behaviors are highlighted by sex separation, female in cyan triangles and male in salmon dots.

**Figure 5:**
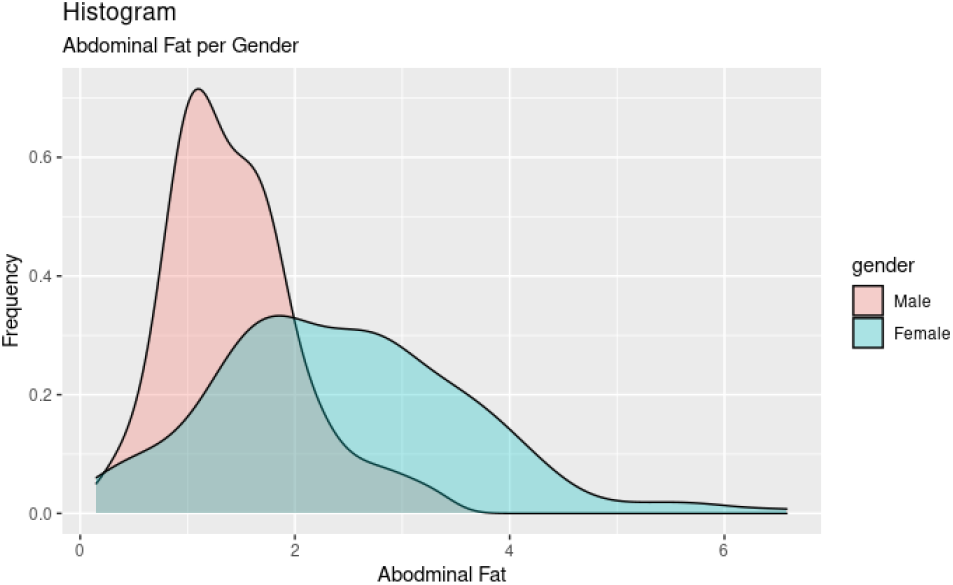
Density plots for abdominal fat content for the male (salmon) and female (cyan) populations of mice.

ANOVAs have similar assumptions, that the variables are independent, the groups have same variance and that residuals are normally distributed. In the case of cholesterol distribution, mainly LDL and VLDL, this is the case. However, with a p-value of 0,320 the null hypothesis cannot be safely disregarded. Figure 6 can provide an intuitive perception as to why this happens.

**Figure 6:**
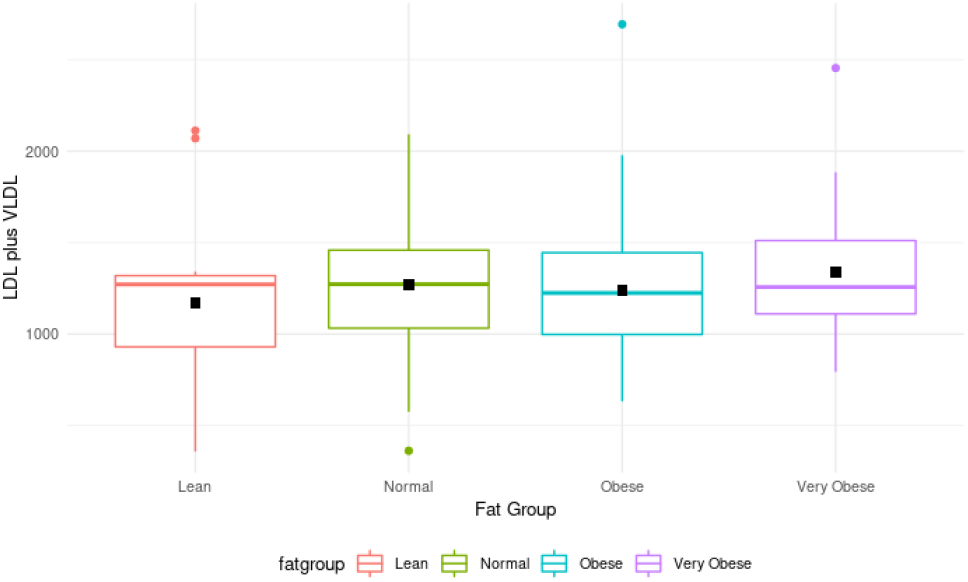
Boxplot of LDL and VLD distributions regarding weight category. The black square represents the location of the mean.

There are, however, various continuous clinical variables that would be interesting to test relating to the weight categories. Two of those variables are leptin and free fatty acids. In order to perform this evaluation MANOVAs provide a suitable statistical tool, however they have many assumptions that need to be verified.

The sample size must have an adequate size. A standard rule of thumb is that the number of variables in each group needs to be bigger than the number of outcome variables, which is the case, as there are 4 outcome variables and the lean group, with least elements, has 12.

Also, there shouldn’t be univariate or multivariate outliers, but there is one extreme outlier in the very obese group.

The data should have univariate and multivariate normality, tested by Shapiro-Wilk test. This data, however does not pass the multivariate normality assumption.

On top of that the data shouldn’t exhibit multicollinearity, which is the case. It is assessed by Pearson’s correlation.

There should be a linearity between all outcome variables for each group. Although it is not perfectly linear due to noisy environment, this assumption is regarded as met.

The covariance matrices should be homogenous, which is tested via rstatix[3] implementation of Box’s M-test, and this assumption is not met.

Finally, the homoscedasticity assumption, tested by Levene’s test also fails for free fatty acids. Considering the number of assumptions that aren’t met for the test, MANOVA would provide unreliable results.

### 2.3 Relation to genetic information

Now, considering the same approach that was taken transforming a continuous data into categories for ANOVA and MANOVA analysis, the same was done regarding the continuous metabolic syndrome variable.

Before applying the factor transformation to analyze differential expression, The sample themselves were subject to the previous described quality check for outliers. This process can be seen on Figure 7. One sample was removed from analysis due to this process

**Figure 7:**
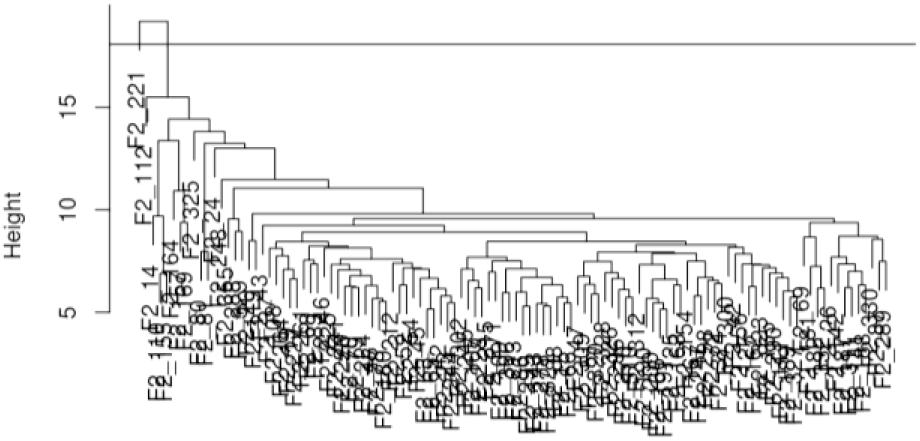
Female samples hierarquical clustering via UMPGA5. The line represents the extreme outlier cutoff height. There is one sample beyond that line, which was discarded.

Instead of performing a graphical analysis for each gene, the assumptions of ANOVA were defined as the conclusions of Shapiro-Wilks and Levene’s test, which were made into a table for all the genes. The table was then filtered considering the p-values of those tests and all then the analysis proceeded as described. A traditional way of observing the differentially expressed genes between two conditions is a volcano plot, which is represented in Figure 8.

**Figure 8:**
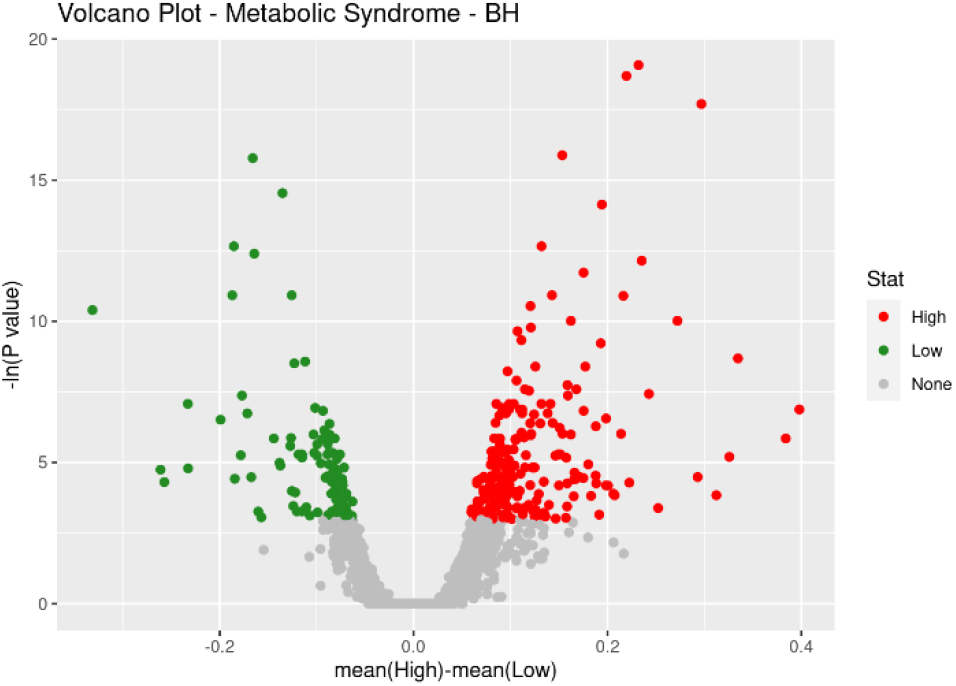
Volcano plot for samples of intense cases of metabolic syndrome vs “control” group. Y axis is -log10 of p-value, so greater values relates to greater significance.

## 4 Discussion

The great diversity of biological data available makes for difficult analysis. In the original clinical information dataset several features were left out of analysis due to their semiquantitative nature and abundant missing values. The best variable of that kind that was maintained, aortic lesions, proved to be unreliable in face of several regression criteria.

On the best model found by the stepwise improvement on aortic lesions, despite not being the preferred metric, it is curious and worth mentioning that the best model has both insulin and glucose/insulin, which, by the scatterplot on Figure 3, have a very defined relationship.

Normally it is not that good to include two variables, one of which dependent on the other, which is why highly correlated variables are removed. That is measured by the Pearson’s correlation coefficient, used in this work. However, Pearson’s coefficient measures the linear dependency between variables and cannot capture this kind of nonlinear relationship. Spearman’s rank coefficient, on the other hand, measures how well the functions can be described by a monotone function and might provide a higher correlation. Nonetheless, given the inclusion of both variables in the best model, one can infer that nonlinear dependence is ignored on the making of a linear model, which means the appropriate correlation coefficient to assay and filter is Pearson’s.

## 5 Conclusions

Analysis of the clinical dataset has shown that there is a significant different metabolic response in males and females. While the molecular metabolic process was not subject to scrutiny, the observed overall phenotype was able to be compared both in an intuitive grasp as seen in the relation between abdominal fat and overall weight, as seen objectively in the t-test, which could differentiate female and male mice according to abdominal fat. This indicates different responses to weight gain and energy storage.

Some variables fare better than others as predictors, and in that sense, aortic lesions was unreliable for the sake of biological conclusions regarding F2 individuals.

Normalization played a vital part allowing features to be to some extent comparable and statistical tests to be done while also allowing for meaningful relations to be extracted.

The method proposed uses ANOVAs for genetic information retrieval and so, it can be used in order to extract genetic meaning from RNA seq or microarray data while accounting for three or more related classes of data. Providing a reliable statistical method to associate genes or clusters of genes with particular traits.

Finally is important to stress that statistical tests each have their own assumption. While they must be met for the sake of consistency, many hypotheses can’t be tested and much data has to be discarded for the sake of analysis.

